# An AI-Driven Decision-Support Tool for Triage of COVID-19 Patients Using Respiratory Microbiome Data

**DOI:** 10.64898/2026.03.18.712739

**Authors:** Eli Gabriel Avina-Bravo, Isabel García-Lorenzo, Mariel Alfaro-Ponce, Luz Breton-Deval

## Abstract

Accurate clinical triage is critical for optimizing decision-making and resource allocation during infectious disease outbreaks such as COVID-19. In this study, we present an AI-driven decision-support tool for the triage of COVID-19 patients based on respiratory microbiome profiles derived from shotgun metagenomic sequencing. We analyzed 477 shotgun respiratory metagenomes from three independent public cohorts and generated genus-level taxonomic profiles, which were integrated with minimal clinical metadata to train supervised machine-learning models, including Random Forest, Support Vector Machine, and XGBoost. Model performance was evaluated using standard classification metrics, cross-validation, and particle swarm optimization for hyperparameter tuning. Across cohorts, we observed a consistent transition from microbiomes dominated by commensal taxa to dysbiotic states enriched in opportunistic and clinically relevant genera, particularly *Acinetobacter* and *Staphylococcus*, in severe and deceased patients. Among the evaluated models, XGBoost consistently achieved the best performance, reaching up to 96.1% accuracy, 97.6% F1-score, and 98.2% ROC–AUC in individual cohorts. When trained on the integrated dataset, XGBoost maintained robust performance (95.1% accuracy, 97.2% F1-score, 94.3% ROC–AUC) and demonstrated greater stability and lower variance compared to alternative models. Feature-importance analyses identified a compact and interpretable set of recurrent microbial predictors, and reduced-feature models retained substantial discriminative power when augmented with key clinical variables. These results support the respiratory microbiome as a valuable source of information for outcome-oriented clinical triage and position microbiome-informed machine learning as a scalable and interpretable decision-support approach for managing COVID-19 and future infectious disease scenarios.

## 1 INTRODUCTION

The coronavirus disease 2019 (COVID-19), caused by the severe acute respiratory syndrome coronavirus 2 (SARS-CoV-2), has posed unprecedented challenges for healthcare systems worldwide, particularly during periods of rapid surge in cases and limited critical-care resources (Schmolke et al., 2025). Five years after the emergence of COVID-19 and nearly two years after the WHO declared the pandemic over (WHO, 2025), the cumulative number of reported cases stands at approximately 778 million. The United States (103 million), China (99.4 million), and India (43 million) report the largest case totals. Globally, reported deaths total about 7.1 million, with the United States, Brazil, and India recording the highest counts; Mexico ranks fifth, following the Russian Federation (WHO, 2025). These data underscore the need to be prepared for the next pandemic.

One of the most critical operational challenges during the pandemic has been the triage of infected patients, as clinical outcomes range widely from asymptomatic or mild disease to severe respiratory failure, multiorgan dysfunction, and death (Negre et al., 2025). Accurate identification of patients at risk of deterioration is therefore essential to optimize clinical decision-making, allocate limited hospital and intensive care resources, and reduce mortality (Khanna et al., 2023; Cervantes-Díaz et al., 2022).

Traditional triage strategies for COVID-19 have relied primarily on clinical assessment, physiological parameters, and selected laboratory biomarkers (Khanna et al., 2023). While these approaches have demonstrated clinical utility, they often lack sufficient sensitivity or specificity for severity prediction in triage settings, particularly during the initial stages of infection when overt clinical deterioration has not yet occurred (Boussen et al., 2022). Moreover, the high heterogeneity of patient responses, influenced by age, comorbidities, immune status, and underlying biological factors, further complicates reliable prognosis using conventional methods alone (Babaiha et al., 2025).

In patients infected with COVID-19, the host microbiome is an endogenous factor that directly influences disease morbidity and mortality; this is especially true when dysbiosis of opportunistic pathogenic bacteria leads them to become pathologically active, resulting in coinfection or superinfection (Mohapatra et al., 2021). Therefore, the study of the respiratory tract microbiome is relevant, since evidence indicates that its composition is a determinant of respiratory health and that its balance may be disrupted by COVID-19 infection (Mohapatra et al., 2021).

Recent advances in high-throughput sequencing technologies have enabled comprehensive characterization of host-associated microbial communities through shotgun metagenomic sequencing (Xie et al., 2024). Metagenomic evidence indicates that the respiratory microbiome undergoes systematic shifts as COVID-19 severity increases (Huang et al., 2024; Shilts et al., 2022; Xie et al., 2024). While uninfected individuals and patients with mild-to-moderate disease display comparable bacterial loads, bacterial burden rises with increasing severity among infected patients (Shilts et al., 2022). Moreover, COVID-19 infection correlates with decreased microbial diversity in the respiratory tract.

However, shotgun metagenomic data are inherently high-dimensional, sparse, and complex, posing substantial challenges for traditional statistical analysis (Bai et al., 2022). Machine learning (ML) techniques offer a powerful framework to address these challenges by automatically extracting informative patterns, interactions, and latent representations from large-scale biological datasets (Khanna et al., 2023). In the context of COVID-19, ML-based models have shown promise in supporting clinical triage, severity prediction, and outcome forecasting using heterogeneous data sources, including clinical variables, imaging, physiological signals, and molecular profiles (Boussen et al., 2022).

Despite these advances, the integration of shotgun metagenomic sequencing data into predictive triage models remains relatively underexplored. Leveraging metagenomic features within ML frameworks has the potential to enhance severity assessment by capturing host–microbiome interactions that precede overt clinical deterioration. Such approaches may provide clinically actionable insights that complement existing triage tools and support data-driven decision-making in both resource-rich and resource-limited healthcare settings (Mitchell et al., 2023).

### 1.1 Motivation

This work introduces and validates an AI-driven, microbiome-informed decision-support framework for clinical triage of COVID-19 patients, demonstrating that respiratory metagenomic signatures, augmented with minimal clinical metadata, can accurately support outcome-oriented patient prioritization across heterogeneous cohorts. Methodologically, the study demonstrates that microbiome-derived features can be effectively leveraged by ensemble learning approaches, particularly XGBoost optimized via particle swarm optimization to achieve robust and generalizable outcome prediction across heterogeneous, multi-center cohorts. Beyond predictive performance, the framework emphasizes interpretability by identifying a compact set of recurrent microbial features that retain high discriminative power, addressing a key limitation of high-dimensional omics-based models in clinical settings.

From a biological perspective, the work provides cross-cohort evidence that COVID-19 severity and mortality are associated with reproducible shifts in respiratory microbial community structure, characterized by the loss of commensal taxa and the expansion of opportunistic and clinically relevant genera such as *Acinetobacter* and *Staphylococcus*. These recurrent microbial signatures, consistently selected by independent models, reflect coherent ecological states rather than dataset-specific noise, reinforcing the biological validity of microbiome-informed severity stratification and supporting the respiratory microbiome as a biomarker of clinical deterioration. Translationally, the study moves beyond descriptive microbiome analysis by proposing a clinically actionable decision-support tool that balances accuracy, robustness, and practicality.

### 1.2 Paper Distribution

In this study, we develop and evaluate supervised ML models for clinical triage of COVID-19 patients using shotgun metagenomic profiles. We systematically compare multiple algorithms and optimization strategies to identify robust predictors of disease severity at different stages. Therefore, this work aims to contribute an AI-driven decision-support framework that integrates metagenomic data to improve clinical triage, resource allocation, and patient outcomes during current and future infectious disease outbreaks. The manuscript is organized as follows: Section 2 describes the clinical datasets, bioinformatic processing pipeline, feature preprocessing, and the machine-learning methodologies employed, including model training, optimization, and evaluation metrics. Section 3 presents the results, beginning with a cross-study comparison of respiratory microbiome profiles, followed by a detailed analysis of model performance, optimization outcomes, and feature importance. Section 4 discusses the implications of the results, emphasizing model robustness, biological interpretability, and clinical relevance, as well as the limitations of the study. Finally, Section 5 concludes the paper by summarizing the main findings and outlining future directions for the integration of microbiome-informed AI tools into clinical decision-support systems.

## 2 MATERIALS AND METHODS

This section details the data acquisition, preprocessing, and computational framework used to construct and evaluate the proposed AI-driven triage models. It describes the clinical cohorts and shotgun metagenomic datasets analyzed, the bioinformatic pipeline used for taxonomic profiling and feature standardization, and the supervised ML algorithms employed for COVID-19 severity classification. Additionally, this section outlines the hyperparameter optimization procedures, cross-validation strategy, and performance metrics used to quantitatively assess model accuracy, robustness, and generalization across heterogeneous datasets.

### 2.1 Clinical Datasets

Globally, multiple groups collected clinical samples during the COVID-19 pandemic to enable in-depth analyses; we identified 14 studies focused on the respiratory microbiome in COVID-19 patients (Gauthier et al., 2022; de Castilhos et al., 2022; Kumar et al., 2022; Hoque et al., 2021; Xie et al., 2024; Feehan et al., 2021; Bai et al., 2022; Rosas-Salazar et al., 2023; Xu et al., 2020; Carvalho et al., 2020; Liu et al., 2021). However, we selected three datasets to train and refine our machine-learning (ML) models: Dataset from PRJCA011099 (Xie et al., 2024) (N=342), from PRJNA743981 (Feehan et al., 2021) (N=98), and from PRJNA781460 (Bai et al., 2022) (N=37). The selected datasets were chosen according to predefined criteria: shotgun metagenomic data, open/public access, and comparable metadata.

It is important to note that the respiratory samples analyzed in this study were collected at different stages throughout patient hospitalization, as defined by the original studies, and do not uniformly correspond to a single standardized timepoint relative to disease onset or admission. Consequently, the present analysis does not assume that all samples represent early-stage infection. Rather, the objective of this work is to leverage heterogeneous microbiome profiles associated with distinct clinical outcomes to train and evaluate supervised machine-learning models capable of learning severity-associated patterns. These learned patterns can subsequently be used to support triage once microbiome data become available in clinical settings. In this context, the proposed framework should be interpreted as a model-training and validation, rather than as a strict evaluation of samples collected exclusively at early disease stages.

### 2.2 Bioinformatic Analysis

All data sets downloaded from NCBI were processed following this pipeline: Raw sequencing reads were quality-filtered, trimmed of sequencing adapters, and subjected to an initial quality-control evaluation using fastp v0.20.1 (Chen et al., 2018; Chen, 2023). fastp was used to remove low-quality reads and adapter contamination and generate a final quality control report for each library. The resulting cleaned reads (hereafter “clean reads”) were carried forward for downstream analysis. Species-level taxonomic profiles were generated from clean reads with MetaPhlAn v4.1.1 (Blanco-Míguez et al., 2023; Truong et al., 2017; Zolfo et al., 2024). The taxonomic abundance table derived from MetaPhlAn was imported into RStudio (Posit team, 2025) and processed using standard microbiome and tidyverse packages (phyloseq, tidyverse, vegan, ggplot2, dplyr, ggpubr).relative-abundance plots were computed and visualized using these packages to characterize community composition and diversity across sample groups. Clean reads were assembled de novo into contigs with MEGAHIT v1.2.9 (Li et al., 2015, 2016)

### 2.3 Data Preprocessing

All models were trained and evaluated on a laptop equipped with an Intel Core i9 processor, 16 GB of RAM, and an NVIDIA RTX 4070 GPU with 8 GB of VRAM. The experiments were implemented in Python (version 3.12) using the following libraries: NumPy Harris et al. (2020)(v1.26), pandas McKinney (2010)(v2.2), scikit-learn Pedregosa et al. (2011)(v1.5), XGBoost Chen and Guestrin (2016)(v2.1), and PySwarms Miranda (2018)(v1.3). Z-score standardization is widely used in ML because it places all features on a common scale and facilitates the detection of outliers or anomalies by emphasizing values that deviate substantially from the mean Sujon et al. (2024); Yerke et al. (2024). Moreover, standardization improves the numerical stability, convergence, and performance of ML models, particularly for algorithms that are sensitive to feature scale or variance. Let **X** ∈ℝ ^*N ×F*^ denote the constructed dataset, where *N* is the number of samples and *F* is the number of features. The standardized value of feature *j* for sample, *i* is defined as

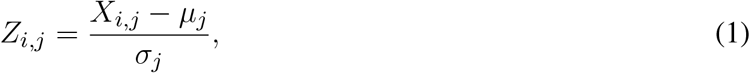

where *X*_*i,j*_ represents the original value of feature *j* in sample *i*, and *µ*_*j*_ and *σ*_*j*_ denote the mean and standard deviation of feature *j*, respectively. This transformation ensures that each feature has a mean of zero and a standard deviation of one.

### 2.4 Machine Learning Techniques

Applying ML techniques to the intelligent triage of COVID-19 patients using shotgun sequencing profiles is crucial for enabling accurate severity assessment to support clinical triaget in a highly complex and high-dimensional biological context (Avila-Ponce de León et al., 2023). Shotgun sequencing generates vast amounts of genomic and metagenomic data that are difficult to interpret using traditional statistical methods alone (Sharpton, 2014). ML models can automatically identify subtle patterns, interactions, and biomarkers within these profiles that are associated with disease progression, immune response, and risk of severe outcomes, allowing clinicians to stratify patients more effectively in the stages of infection (Ng et al., 2023).

Moreover, ML–driven triage supports faster, data-driven clinical decision-making, which is essential during pandemics where healthcare resources are limited. By predicting patient severity early, these models help prioritize critical care (Pundkar et al., 2025), optimize hospital resource allocation, and reduce mortality rates. Integrating ML with sequencing data also enables continuous model improvement as new data become available, making the triage system adaptive, scalable, and more resilient to emerging variants or population-specific responses.

In this framework, each patient is represented by a feature vector **x**_*i*_ ∈ ℝ^*d*^ extracted from shotgun sequencing profiles, where *d* denotes the number of metagenomic features, including microbial abundances, functional gene counts, and derived descriptors. These feature representations are used consistently across all ML models evaluated in this study to support COVID-19 severity prediction.

#### 2.4.1 Random Forest

Random Forest (RF) is a ensemble learning method composed of *T* decision trees, each trained on a bootstrap sample of the patient dataset. For COVID-19 severity classification, each tree maps the input feature vector to a severity class, and the final prediction is obtained by majority voting:

For COVID-19 severity classification, each tree maps the sequencing-derived features to a severity class *c* ∈ 𝒞. The final prediction is obtained by majority voting:

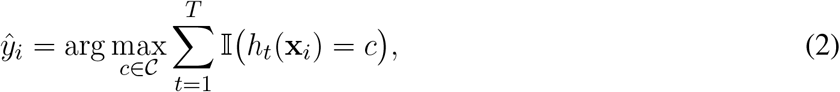

where 𝕀 (*·*) is the indicator function.

At each node, a random subset of metagenomic features ℱ_*t*_ *⊂*{1, …, *d*} is evaluated to determine the optimal split, promoting model diversity and robustness to high-dimensional and sparse shotgun sequencing data.

#### 2.4.2 Support Vector Machines

Support Vector Machines (SVM) aim to learn an optimal decision function that separates patient severity classes with maximum margin in a high-dimensional feature space. The binary classification problem is formulated as the following constrained optimization task:

For the binary classification case, the SVM solves the following optimization problem:

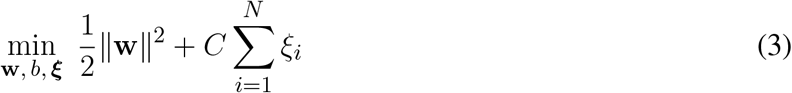

subject to

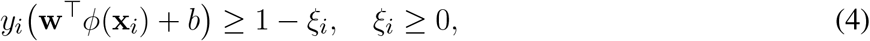

where *y*_*i*_ ∈ {*−*1, +1}denotes the severity label, *ϕ*(*·*) is a kernel-induced feature mapping, *ξ*_*i*_ are slack variables allowing misclassification, and *C* is a regularization parameter.

The resulting decision function is given by:

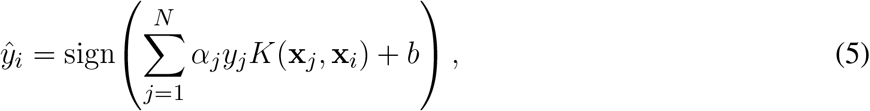

#### 2.4.3 XGBoost

XGBoost models COVID-19 severity prediction as an additive ensemble of regression trees *T*, optimized through gradient boosting and regularization to capture non-linear interactions among metagenomic features.

The predicted output is given by:

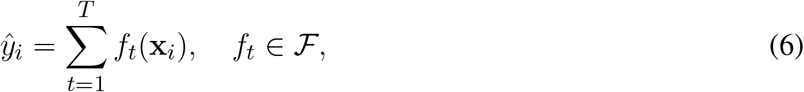

where ℱ is the space of decision trees. The model is trained by minimizing a regularized objective function:

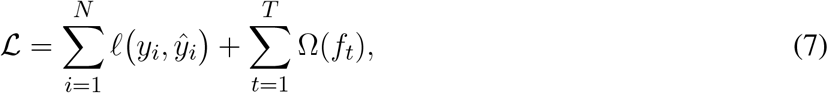

with 𝓁 (*·*) denoting a differentiable loss function (e.g., logistic loss for severity classification) and Ω(*f*_*t*_) a regularization term that controls model complexity:

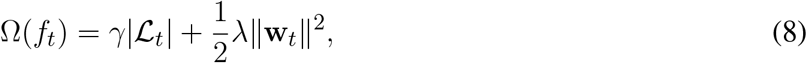

where |ℒ_*t*_| is the number of leaves in tree *t*, **w**_*t*_ are the leaf weights, and *γ* and *λ* are regularization parameters.

By iteratively fitting trees to the gradients of the loss function, XGBoost efficiently captures non-linear interactions and hierarchical patterns in high-dimensional and sparse shotgun sequencing data, making it well suited for COVID-19 severity assessment for clinical triage.

### 2.5 Model Training and Optimization

Model training in the context of COVID-19 severity assessment focuses on enabling ML models to identify discriminative patterns from shotgun sequencing profiles related to disease progression. Model parameters are optimized using labeled patient data by minimizing a prediction error loss function. Common techniques include tree-based methods such as RF and XGBoost, margin-based optimization in SVM, gradient-based optimization, regularization strategies to prevent overfitting, and K-fold cross-validation for robust model selection and hyperparameter tuning in limited and heterogeneous sequencing datasets.

Particle Swarm Optimization (PSO) is employed as a population-based metaheuristic optimization technique to tune model hyperparameters and improve predictive performance. In PSO, a swarm of *P* particles explores the search space, where each particle represents a candidate solution (e.g., a set of model hyperparameters). The position 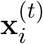and velocity 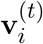of particle *i* at iteration *t* are updated according to:

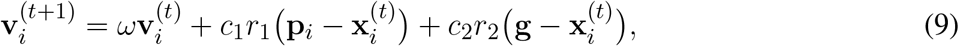

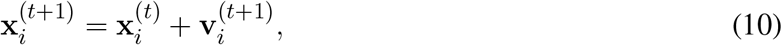

where **p**_*i*_ is the best position previously found by particle *i*, **g** is the global best solution found by the swarm, *ω* is the inertia weight controlling exploration, *c*_1_ and *c*_2_ are acceleration coefficients, and *r*_1_, *r*_2_*∼* 𝒰 (0, 1) are random variables. The fitness of each particle is evaluated using a performance metric derived from COVID-19 severity prediction, allowing PSO to efficiently search for optimal hyperparameter configurations in complex and high-dimensional spaces.

K-fold cross-validation is employed to robustly evaluate model generalization performance when predicting COVID-19 severity from shotgun sequencing data. The dataset is partitioned into *K* approximately equal-sized folds; at each iteration, one fold is used for validation while the remaining *K−* 1 folds are used for training. This procedure is repeated *K* times so that each patient sample contributes to both training and validation. The overall cross-validation performance is estimated as:

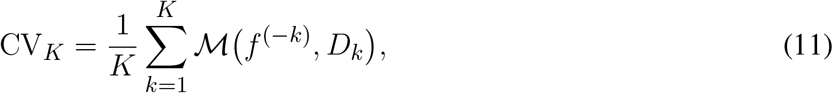

where *f* ^(*−k*)^ denotes the model trained on all folds except the *k*-th fold, *D*_*k*_ represents the validation dataset for fold *k*, and ℳ(*·*) is the evaluation metric used to quantify COVID-19 severity prediction performance.

#### 2.5.1 Performance Metrics

To evaluate the performance of ML models for COVID-19 severity assessment using shotgun sequencing profiles, standard classification metrics including Accuracy, Precision, Recall, and F1-score are employed. These metrics are computed from the confusion matrix, which summarizes model predictions in terms of true positives (TP), true negatives (TN), false positives (FP), and false negatives (FN). In this context, TP represents the number of severe COVID-19 patients correctly classified as severe, TN corresponds to mild patients correctly classified as mild, FP denotes mild patients incorrectly predicted as severe, and FN represents severe patients incorrectly predicted as mild.

Accuracy measures the overall proportion of correctly classified patients and is defined as:

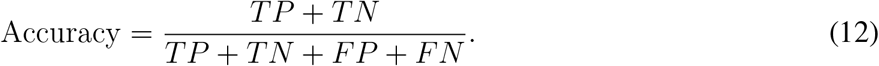

Precision evaluates the reliability of severe case predictions, indicating the proportion of correctly identified severe patients among all patients predicted as severe:

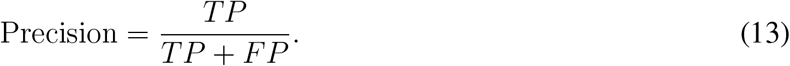

Recall (also referred to as sensitivity) measures the ability of the model to correctly identify patients with severe COVID-19:

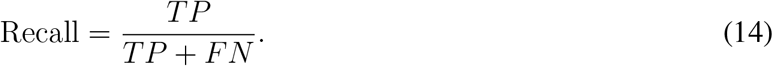

The F1-score provides a harmonic balance between Precision and Recall, which is particularly important for imbalanced clinical datasets, and is defined as:

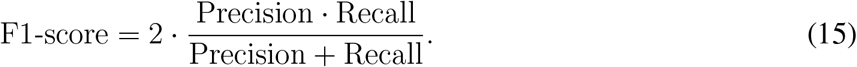

The Receiver Operating Characteristic Area Under the Curve (ROC AUC) is defined as the area under the ROC curve, which plots the Precision against the Recall:

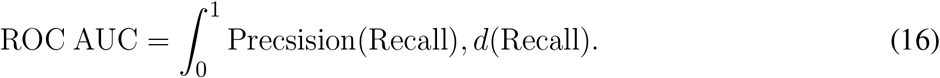

ROC AUC values range from 0 to 1, where a value of 0.5 indicates no discriminative ability (equivalent to random classification), and a value of 1.0 indicates perfect discrimination. Unlike Accuracy and threshold-dependent metrics, ROC AUC provides a global assessment of model performance and is particularly suitable for imbalanced clinical datasets. Together, these metrics provide a comprehensive and clinically relevant evaluation of model performance for COVID-19 severity prediction based on shotgun sequencing data.

### 2.6 Datasets Characteristics

Across the different datasets, a variety of metadata variables were identified. However, because the purpose of this work is to correlate multiple datasets in order to improve ML models, only metadata that were consistently reported across all datasets were analyzed. As shown in Table 1, these variables include age, gender, history of antibiotic treatment, COVID-19 test positivity, and disease severity. Although additional metadata could potentially improve model performance by providing richer clinical context for each sample, such information was not uniformly available across studies and was therefore excluded to avoid introducing bias.

**Table 1.** Consistent metadata found on the datasets.

| Dataset | Age | Gender (Female) | Antibiotic (Yes) | Severity | COVID-19 (Positive) | Outcome (Alive) |
| --- | --- | --- | --- | --- | --- | --- |
| PRJCA011099 | 58.21±19.44 | 47.66% | 93.27% | Severe-55.55%<br>Mild-44.44% | 100% | 82.45% |
| PRJNA743981 | 51.42±16.48 | 66.32% | 17.34% | Light-2.04%<br>Very Mild-20.40%<br>Mild-43.87%<br>Moderate-19.38%<br>Severe-12.24%<br>High-2.07% | 79.59% | 94.89% |
| PRJNA781460 | 59.32±9.93 | 18.91% | 16.21% | NA | 78.37% | 78.37% |

The mean age of participants is comparable across datasets, ranging from approximately 51 to 59 years, with Datasets PRJCA011099 and PRJNA781460 including older populations on average. Gender distribution varies substantially across studies: Dataset PRJNA743981 shows the highest proportion of female participants (66.32%), whereas Dataset PRJNA781460 is predominantly male. The prevalence of antibiotic treatment also differs markedly, being very high in Dataset PRJCA011099 (93.27%) and considerably lower in Datasets PRJNA743981 and PRJNA781460.

Disease severity distributions further highlight inter-dataset heterogeneity. Dataset PRJCA011099 is skewed toward severe cases, while Dataset PRJNA743981 spans a broad spectrum of severity levels, from very mild to high severity, indicating greater clinical diversity. Severity information is not available for Dataset PRJNA781460. COVID-19 test positivity rates remain relatively high across datasets; however, Datasets PRJNA743981 and PRJNA781460 include subsets of participants without confirmed positive tests.

Clinical outcomes also vary across studies. Dataset PRJNA743981 reports the highest proportion of surviving patients (94.89%), followed by Dataset PRJCA011099 (82.45%) and Dataset PRJNA781460 (78.37%).

From each study, shotgun metagenomic data were retrieved on a per-sample basis and analyzed (see Section 2.2), from which sequencing read counts and relative abundances were computed at the genus level.

The total number of sequencing reads (SR) for sample *j* is defined as

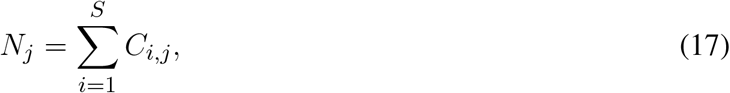

where *S* denotes the total number of genus, *C*_*i,j*_ represents the number of reads mapped to genus *i* in sample *j*, and *j* indexes the samples. The relative abundance (RA) of genus *i* in sample *j* is defined as the proportion of reads assigned to that genus relative to the total number of reads in the sample:

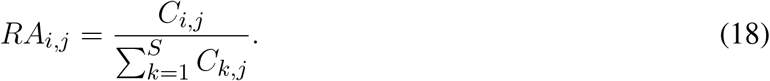

This procedure results in a table reporting the sequencing reads (SR) and relative abundances (RA) of individual bacterial genera for each sample. However, this representation is not directly suitable for ML applications, as the set of detected bacterial genera varies across samples within a dataset. To standardize the feature space, the table was transformed so that each sample contains the same set of bacterial genera. Genera not detected in a given sample were assigned zero values for both SR and RA.

This transformation inevitably produces a sparse matrix, with a large proportion of zero-valued entries. To mitigate sparsity and reduce noise, only bacterial genera detected in at least 10 samples were retained for subsequent analysis. Table 2 presents the resulting set of bacterial genera identified in each dataset. Genera that are shared across multiple datasets are highlighted in blue.

**Table 2.**
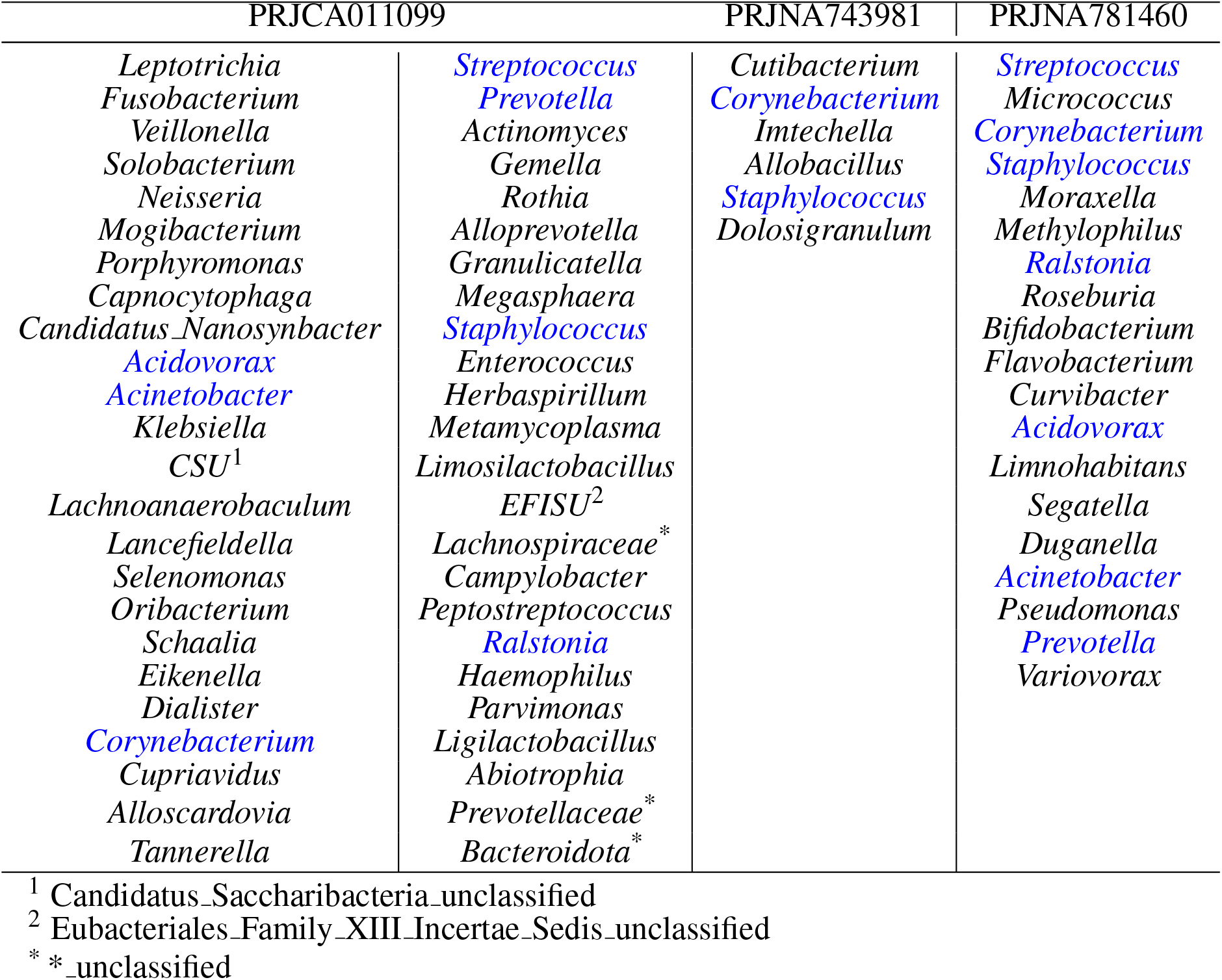
Resulting bacterial genes identified in each dataset.

Dataset PRJCA011099 exhibits the greatest taxonomic diversity, containing a broad range of oral- and respiratory-associated genera, whereas Datasets PRJCA011099 and PRJNA781460 show more restricted genus sets. Several bacterial genera, including *Streptococcus, Corynebacterium, Staphylococcus, Prevotella, Acinetobacter, Ralstonia*, and *Acidovorax*, are consistently observed across datasets, suggesting a core group of taxa common to multiple studies. In contrast, a substantial number of genera are unique to individual datasets, reflecting differences in cohort characteristics, sampling strategies, and sequencing or preprocessing pipelines.

## 3 RESULTS

### 3.1 Cross-study Comparison of Respiratory Tract Microbiome Profiles

Across the three datasets, a consistent pattern of respiratory microbiome was observed along the severity gradient, characterized by the progressive loss of commensal taxa and the expansion of opportunistic and hospital-associated genera (see Figure **??**). In Dataset PRJCA011099, healthy individuals were dominated by oral-associated genera such as *Prevotella, Veillonella, Streptococcus, Capnocytophaga*, and *Gemella*. During mild COVID-19, pronounced blooms of *Streptococcus* and *Gemella* were observed, accompanied by reductions in *Prevotella* and *Capnocytophaga*. As severity increased, low-abundance taxa expanded markedly, including *Acinetobacter, Metamycoplasma*, and *Herbaspirillum*, while several early-dominant genera declined. In deceased patients, the microbial profile was strongly dominated by *Acinetobacter* and *Staphylococcus*, alongside a collapse of multiple commensal taxa.

A similar transition was evident in dataset PRJNA743981, despite its distinct baseline composition. Healthy individuals exhibited a microbiome dominated by *Corynebacterium* and *Cutibacterium*, with minor contributions from oral-associated genera. Mild and moderate disease was associated with decreases in *Corynebacterium, Imtechella*, and *Allobacillus*, and concurrent increases in *Cutibacterium, Staphylococcus, Dolosigranulum*, and *Moraxella*. In severe cases, this pattern inverted: several genera that expanded in mild disease declined sharply, while *Corynebacterium, Streptococcus*, and *Alloprevotella* increased, indicating a shift toward a more dysbiotic and clinically associated community structure.

In dataset PRJNA781460, where severity was defined using oxygen saturation, non-severe cases were dominated by *Corynebacterium* and *Cutibacterium*, whereas severe cases exhibited enrichment of *Flavobacterium* and *Staphylococcus*, along with reductions in *Cutibacterium, Dolosigranulum*, and *Streptococcus*. Several genera detected in non-severe patients were absent in severe cases, reinforcing the pattern of community simplification observed across datasets.

Together, these results demonstrate that, despite differences in baseline community composition and severity definitions, all three datasets converge on a shared ecological trajectory: COVID-19 is associated with selective blooms of specific taxa, while severe disease is characterized by the loss of commensal diversity and the dominance of opportunistic genera. This consistency across cohorts supports the robustness of severity-associated microbiome patterns and provides a biological foundation for the predictive modeling framework.

Preliminary analyses indicated that the features Disease Severity and COVID-19 test status introduced bias into the models, as both variables are strongly correlated with patient outcomes. Consequently, these features were excluded from the analysis, and the results reported below were obtained using the reduced feature set.

As shown in Table 3, XGBoost (XGB) outperformed the other models in Datasets PRJCA011099 and PRJNA743981, while RF achieved slightly better performance in Dataset PRJNA781460. Overall, XGB demonstrates consistently strong performance across datasets, particularly in terms of F1-score and ROC–AUC. For Datasets PRJCA011099 and PRJNA743981, F1-score and ROC–AUC values exceed 97%, indicating an excellent balance between precision and recall and a near-perfect ability to discriminate clinical outcomes. These results suggest that the model achieves high predictive accuracy while effectively minimizing both false positives and false negatives. Performance on Dataset PRJNA781460 is comparatively lower for all models, likely reflecting the smaller sample size and increased variability in this dataset.

**Table 3.** Model’s performance across the three datasets.

| Datasets | Models | Accuracy | Precision | Recall | F1-score | ROC-AUC |
| --- | --- | --- | --- | --- | --- | --- |
| PRJCA011099 | SVM | 83.49% | 93.58% | 85.88% | 89.57% | 87.71% |
|  | RF | 93.20% | 96.42% | 95.29% | 95.85% | 97.64% |
|  | XGB | 96.11% | 98.79% | 96.47% | 97.61% | 98.16% |
| PRJNA743981 | SVM | 93.33% | 93.33% | 100% | 96.55% | 80.35% |
|  | RF | 93.33% | 93.33% | 100% | 96.55% | 94.64% |
|  | XGB | 96.66% | 96.55% | 100% | 98.24% | 94.64% |
| PRJNA781460 | SVM | 58.33% | 75.00 % | 66.66% | 70.58% | 55.55% |
|  | RF | 75.00% | 75.00% | 100% | 85.71% | 55.55% |
|  | XGB | 75.00% | 80.00% | 88.88% | 84.21% | 51.85% |

Table 4 reports the performance of models trained on the combined dataset obtained by merging Datasets PRJCA011099, PRJNA743981, and PRJNA781460. This integration increases the total number of samples to 477, thereby providing the ML models with greater variability and a broader representation of clinical patterns. Although overall performance metrics are lower than those obtained when training on Datasets PRJCA011099 and PRJNA743981 individually, XGB remains the best-performing model across all evaluated metrics.

**Table 4.** Model’s performance on the three dataset combined.

| Models | Accuracy | Precision | Recall | F1-score | ROC-AUC |
| --- | --- | --- | --- | --- | --- |
| SVM | 85.41% | 92.43 % | 90.16% | 91.28% | 78.98% |
| RF | 90.97% | 92.91% | 96.72% | 94.77% | 93.01% |
| XGB | 93.05% | 93.75% | 98.36% | 96.00% | 93.10% |

The observed decrease in performance is likely attributable to the inclusion of Dataset PRJNA781460, which contains a higher proportion of patients with diseased clinical outcome. This imbalance introduces additional heterogeneity and increases the number of possible feature combinations associated with the outcome, making classification more challenging. Nevertheless, XGB maintains strong discriminative capability, achieving an F1-score of 96.00% and a ROC–AUC of 93.10%, indicating robust performance even under increased dataset heterogeneity.

Considering these results, the two best-performing models (RF and XGB) were further optimized using the Particle Swarm Optimization (PSO) algorithm (see Section 2.5). As both models are based on decision-tree ensembles, they share several key hyperparameters, most notably the maximum tree depth and the number of estimators. Consequently, identical search ranges were defined for these parameters in both models. In contrast, the learning rate was optimized only for XGB, as this parameter is not applicable to RF.

Table 5 summarizes the PSO configuration and the hyperparameter search space explored for each model. The maximum tree depth was varied over a wide range to capture both shallow and deep tree structures, while the number of estimators was constrained to balance model expressiveness and computational cost.

**Table 5.** Model’s hyperparameters to be optimize using this PSO configuration.

| Models | Max Depth | Num Estimators | Learning Rate | $c_1$ | $c_2$ | $\omega$ | $P$ |
| --- | --- | --- | --- | --- | --- | --- | --- |
| RF | [5 - 250] | [50 - 200] | NA | 2 | 2 | 0.75 | 30 |
| XGB |  |  | [0.1 - 0.5] |  |  |  | 45 |

For PSO, identical cognitive and social coefficients (*c*_1_ = *c*_2_ = 2) and inertia weight (*ω* = 0.75) were used for both models to ensure a fair optimization framework. A larger swarm size (i.e., *P*) was employed for XGB compared to RF, reflecting the higher dimensionality of its hyperparameter space due to the inclusion of the learning rate.

Each model was optimize during 100 epochs, using 1*−* F1-score as the PSO cost function. Table 6 reports the performance obtained after optimization. When compared with the results in Table 4, an overall improvement in performance is observed, particularly for XGB, where both the F1-score and ROC-AUC increased by approximately 1.2%. Although RF also exhibited improvements in most evaluation metrics, its ROC-AUC decreased after optimization. This suggests that, despite higher point-wise classification performance, the model struggles to consistently discriminate between false positives and false negatives across different decision thresholds.

**Table 6.** Performance of the optimized models using PSO.

| Models | Accuracy | Precision | Recall | F1-score | ROC-AUC |
| --- | --- | --- | --- | --- | --- |
| RF | 91.66% | 94.35% | 95.90% | 95.12% | 89.79% |
| XGB | 95.13% | 94.57% | 100% | 97.21% | 94.26% |

Using the PSO-optimized hyperparameters, a 5-fold cross-validation procedure was performed for both models. As shown in Table 7, the XGB model consistently outperforms the RF model across all evaluation metrics, although the performance margin remains modest. Notably, XGB exhibits lower standard deviations across folds, indicating greater stability and robustness to data partitioning. Moreover, this reduced variability suggests improved generalization and more reliable predictive behavior across different subsets of the data.

**Table 7.** Cross-Validation Results.

| Models | Accuracy | Precision | Recall | F1-score | ROC-AUC |
| --- | --- | --- | --- | --- | --- |
| RF | 90.35±03.22% | 92.78±03.57% | 96.06±01.36% | 94.35±02.03% | 91.58±04.98% |
| XGB | 91.61±02.22% | 93.50±02.90% | 96.81±01.86% | 95.09±01.47% | 92.86±04.25% |

One of the main objectives in this work is to analyze and find discriminating factors that healthcare personnel could use to triage COVID-19 patients. From the trained models, the top 30 features were analyzed in Table 8. To each feature was associated an importance value that represents the weight in each model that contributes to the classification result.

**Table 8.**
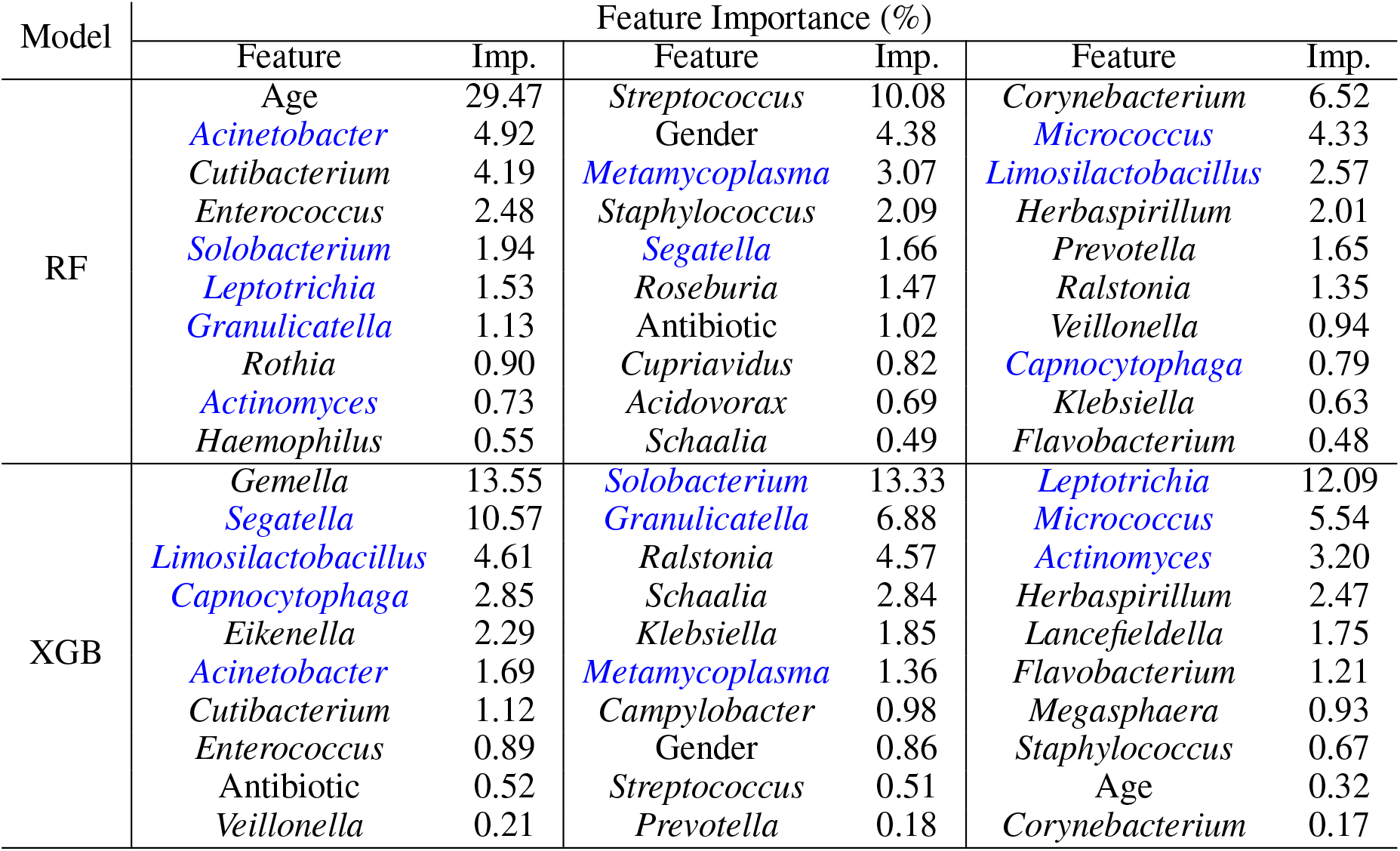
Top-ranked feature importance values obtained from Random Forest and XGBoost models.

To identify the top 10 features associated to both models, the Algorithm 1 was used. Recurrent features were defined as predictors assigned non-zero importance by both RF and XGBoost models, reflecting model-consistent relevance rather than rank-dependent importance. From the list obtained, the most recurrent features were highlighted in blue in Table 8.

#### Algorithm 1

Importance-Based Identification of Recurrent Features

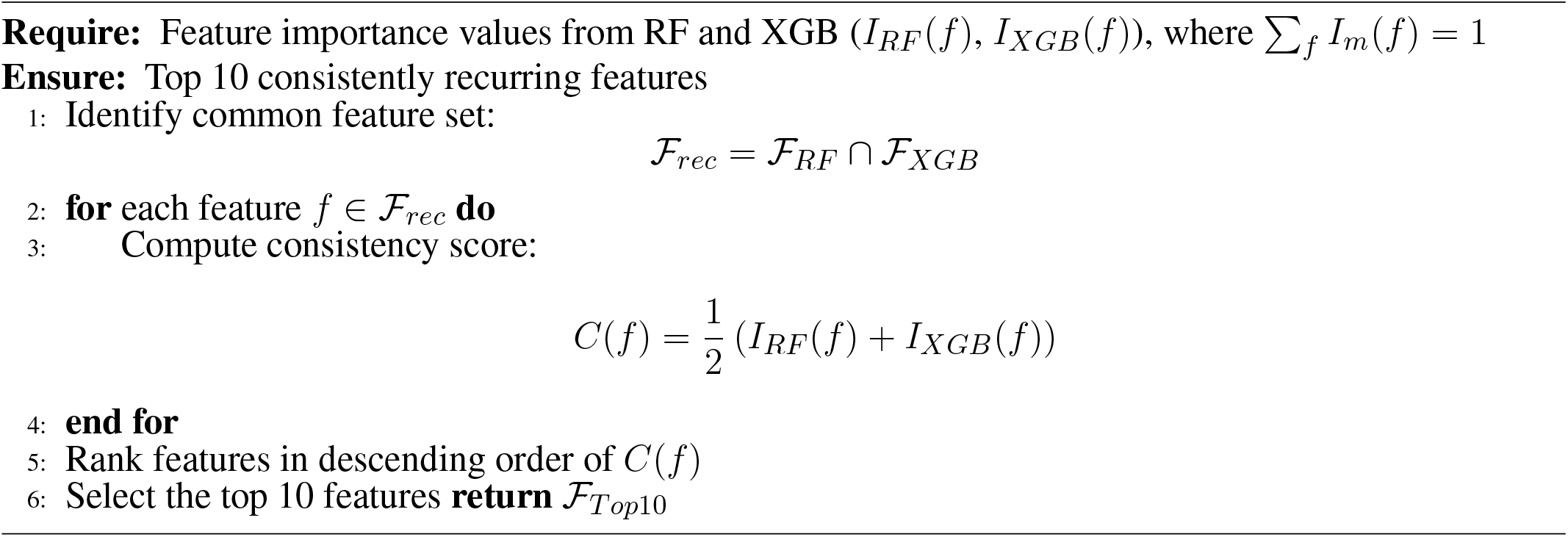

As a final validation step, the subset of highlighted features was used to assess whether the models were robust enough to correctly classify patient outcomes using only a reduced set of predictors. Specifically, the RF and XGB models were retrained using only the top 10 recurrent features identified across both models. Default hyperparameters were employed for this experiment, as the PSO-optimized hyperparameters were tuned for the full feature space and are therefore not directly applicable to a reduced feature set.

The results of this experiment are reported in Table 9. When only the top 10 recurrent features were used, a noticeable degradation in performance was observed for both models compared to the results obtained with the full feature set (see Table 4). On average, the reduction in performance across most metrics was approximately 5%, with the exception of Recall and ROC-AUC. While Recall remained largely stable—indicating that the models retained their ability to correctly identify positive cases—the substantial decrease in ROC-AUC suggests a reduced capability to distinguish between false positives and false negatives across varying classification thresholds.

**Table 9.** Results with only the selected features.

| Models | Accuracy | Precision | Recall | F1-score | ROC-AUC |
| --- | --- | --- | --- | --- | --- |
| Using the top 10 recurrent features |  |  |  |  |  |
| RF | 84.72% | 87.87% | 95.08% | 91.33% | 78.68% |
| XGB | 88.19% | 88.88% | 98.36% | 93.38% | 78.05% |
| Using the top 10 recurrent features with also Age and *Gemella*. |  |  |  |  |  |
| RF | 92.36% | 94.39% | 96.72% | 95.54% | 92.88% |
| XGB | 93.75% | 95.93% | 96.72% | 96.32% | 95.67% |

To further investigate this behavior, the top-ranked feature from each model (i.e., Age for RF and *Gemella* for XGB; see Table 8) was added to the set of top 10 recurrent features. Under this configuration, both models exhibited a marked improvement in performance. In particular, the RF model achieved results approximately 1% higher than those reported in Table 6, while the XGB model showed competitive, albeit slightly lower, performance. Although these results are not directly comparable to those obtained using PSO-optimized hyperparameters, they nonetheless outperform the baseline models trained on the full feature set without optimization (see Table 4).

Overall, these findings highlight that a compact and interpretable subset of features—when augmented with the most influential individual predictors—can retain a large portion of the predictive power of the full model, while offering improved practicality for clinical decision support.

### 3.2 Biological Patterns Underlying Model-Selected Microbial Features

Analysis of feature importance across models trained on the integrated dataset revealed a recurrent set of microbial genera consistently associated with COVID-19 severity (see Table 8). These genera were not randomly distributed but clustered into ecologically coherent groups. Features enriched among severe patients were dominated by taxa commonly detected in respiratory dysbiosis and clinical settings, including *Acinetobacter, Staphylococcus, Klebsiella, Enterococcus*, and *Corynebacterium*. In contrast, features associated with non-severe cases included genera typically abundant in healthy upper respiratory communities, such as *Prevotella, Veillonella, Capnocytophaga, Gemella*, and *Leptotrichia*. This structured pattern indicates that the model captures consistent shifts in community composition rather than dataset-specific noise. In particular, these recurrent microbial signatures were identified from models trained in the combined data set comprising all three cohorts, supporting their robustness in heterogeneous sampling strategies and patient populations. The convergence of features selected by the model with biologically coherent community profiles provides evidence that the predictive framework reflects the underlying structure of the microbial community associated with disease progression, reinforcing the biological validity of microbiome-informed severity stratification.

## 4 DISCUSSION

This study demonstrates that respiratory microbiome profiles can be effectively leveraged within an AI-driven framework to support clinical triage of COVID-19 patients. While previous microbiome studies in COVID-19 have primarily focused on descriptive community shifts or retrospective associations, the results presented here show that microbiome-derived features can be transformed into robust, generalizable, and clinically actionable decision-support signals. By integrating shotgun metagenomic data from three independent cohorts, the proposed framework moves beyond single-cohort observations and addresses a key limitation of prior studies: limited generalizability across heterogeneous patient populations. Although samples were collected at heterogeneous stages during hospitalization, the consistent severity-associated microbial patterns identified across cohorts enable the training of models intended to support triage once sequencing data are obtained.

A central finding of this work is the consistent predictive performance of ensemble learning models—particularly XGBoost—across datasets with distinct baseline microbiome compositions and severity definitions. In individual cohorts, XGBoost achieved F1-scores and ROC–AUC values exceeding 97%, and importantly, maintained strong performance when trained on the integrated dataset (n = 477), despite increased heterogeneity. This stability, further supported by lower variance in cross-validation, highlights the suitability of gradient-boosted decision trees for microbiome-based triage tasks, where feature interactions are non-linear and data sparsity is common. These results directly support the manuscript’s contribution as an AI-driven decision-support tool rather than a cohort-specific predictive model.

Beyond predictive accuracy, the analysis of feature importance revealed that model performance is driven by recurrent and biologically coherent microbial signatures, rather than idiosyncratic features unique to individual datasets. Across RF and XGBoost models, severe outcomes were consistently associated with enrichment of opportunistic and clinically relevant genera such as *Acinetobacter, Staphylococcus, Klebsiella*, and *Enterococcus*, whereas non-severe outcomes were characterized by commensal taxa including *Prevotella, Veillonella, Gemella*, and *Leptotrichia*. The convergence of these features across models and cohorts indicates that the framework captures stable ecological states linked to disease progression, reinforcing the biological validity of microbiome-informed severity stratification. This cross-model and cross-cohort consistency represents a key contribution of the study.

Importantly, the study shows that effective triage does not require high-dimensional feature spaces. Models trained using a reduced set of recurrent microbial predictors retained much of the predictive power of the full models, and performance was largely recovered when combined with minimal clinical context, particularly age. This finding is critical from a translational standpoint, as it demonstrates that compact and interpretable models can support decision without sacrificing accuracy, addressing a major barrier to the clinical adoption of omics-based AI tools. Rather than acting as black-box predictors, the proposed models provide transparent decision support grounded in biologically interpretable features.

Taken together, the results position the respiratory microbiome as a viable biomarker for COVID-19 severity and establish the feasibility of AI-driven, microbiome-informed decision-support systems for clinical triage. By emphasizing robustness, interpretability, and reduced feature dependence, this work contributes a scalable framework that can be adapted to future infectious disease outbreaks, where rapid and reliable risk stratification is essential for optimizing healthcare resource allocation.

Despite the strong and consistent performance observed across cohorts, several limitations should be acknowledged. First, this study relies on retrospective, publicly available datasets, which limits control over sampling protocols, sequencing depth, and clinical workflows, highlighting the need for prospective validation in real-world hospital settings. Second, the models operate on single timepoint microbiome profiles and therefore do not capture temporal dynamics of microbial changes during disease progression. Third, clinical metadata availability was limited and heterogeneous across cohorts, restricting the integration of additional variables such as comorbidities or laboratory markers that may further enhance predictive performance. Antibiotic exposure, while included as a model feature, remains a potential confounder given its known impact on respiratory microbial communities. Finally, analyses were conducted at the genus level, which may obscure species- or strain-specific effects relevant to pathogenesis. Addressing these limitations through longitudinal sampling, richer clinical annotation, and prospective clinical validation will be essential for translating microbiome-informed AI triage tools into routine clinical practice.

## 5 CONCLUSION

This study demonstrates that respiratory microbiome profiles derived from shotgun metagenomic sequencing can be systematically integrated into an AI-driven decision-support framework for the clinical triage of COVID-19 patients. Using data from three independent cohorts and a combined dataset comprising 477 samples, ensemble machine-learning models, particularly XGBoost, consistently achieved high discriminative performance, with F1-scores exceeding 97% in individual datasets and remaining above 96% when trained on the integrated cohort. The relatively small reduction in performance observed under increased heterogeneity, together with reduced variance in cross-validation, indicates that gradient-boosted decision trees exhibit stable generalization behavior when exposed to diverse microbiome compositions and severity definitions.

Analysis of model outputs revealed that predictive performance is driven by recurrent and biologically structured feature patterns rather than dataset-specific artifacts. Feature-importance analyses showed that severe and deceased outcomes are consistently associated with increased contributions from opportunistic and clinically relevant genera such as *Acinetobacter, Staphylococcus, Klebsiella*, and *Enterococcus*, whereas non-severe outcomes are characterized by higher importance assigned to commensal taxa including *Prevotella, Veillonella, Gemella*, and *Leptotrichia*. This convergence across RF and XGBoost models indicates that the learned decision boundaries reflect coherent microbiome states linked to disease progression, supporting the biological plausibility of the decision-support framework.

From a modeling perspective, experiments using reduced feature sets further highlight the trade-off between dimensionality and discriminative power. While restricting the models to the top recurrent microbial features led to moderate decreases in ROC–AUC, the recovery of performance upon inclusion of key clinical variable, particularly age, demonstrates that compact and interpretable feature representations can preserve much of the predictive signal. This behavior is consistent with ensemble models leveraging non-linear interactions between microbiome composition and host-related factors to support triage decisions.

Overall, the results indicate that microbiome-informed machine learning can provide robust, interpretable, and scalable decision support for clinical triage, rather than functioning as a cohort-specific predictive model. Although prospective validation and longitudinal analyses are required prior to deployment, the observed stability, feature consistency, and performance under heterogeneous conditions establish a strong methodological and biological foundation for incorporating respiratory metagenomic data into data-driven triage systems for COVID-19 and future infectious disease scenarios.

**Figure 1.**
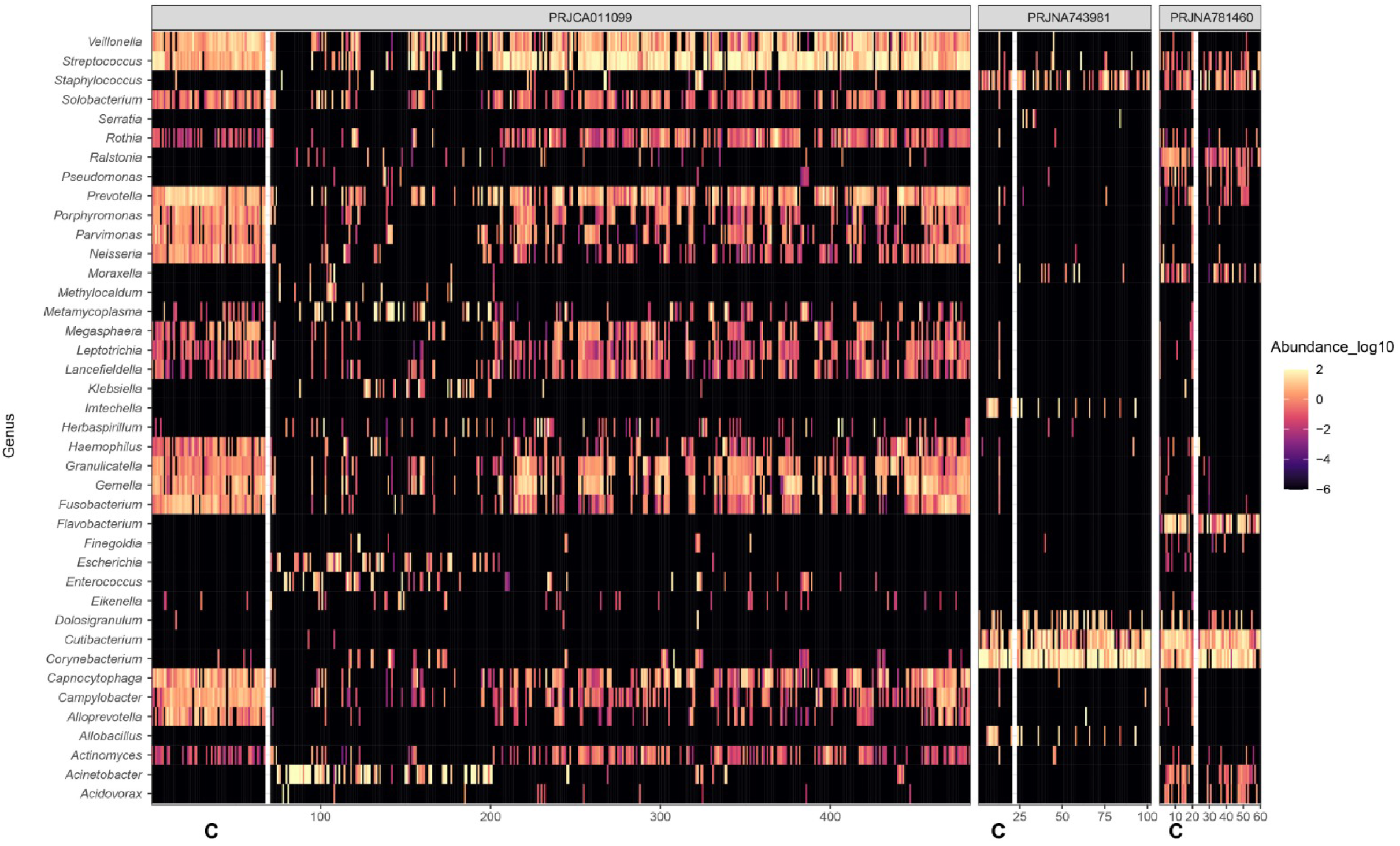
Respiratory microbiome profiles in datasets. Where C denotes control patients.

## Ethics statements

### Studies involving animal subjects

Generated Statement: No animal studies are presented in this manuscript.

### Studies involving human subjects

Generated Statement: No human studies are presented in the manuscript.

## Inclusion of identifiable human data

Generated Statement: No potentially identifiable images or data are presented in this study.

## Data availability statement

Generated Statement: The datasets presented in this study can be found in online repositories. The names of the repository/repositories and accession number(s) can be found in the article/supplementary material.

## Generative AI disclosure

No Generative AI was used in the preparation of this manuscript.

## AUTHOR CONTRIBUTIONS

EGAB: Conceptualization, Methodology, Formal analysis, Visualization, Writing – original draft, Validation. IGL: Data curation, Methodology, Writing – original draft, Investigation. MAP: Writing – original draft, Data curation, Conceptualization, Writing – review & editing, Formal analysis. LBD: Writing – review & editing, Conceptualization, Investigation, Methodology, Formal analysis, Project administration.

## ACKNOWLEDGMENTS

Special thanks to Juan Manuel Hurtado Ramirez for his assistance with IT support.

